# Disrupted Energy Landscape in Individuals with Mild Cognitive Impairment: Insights from Network Control Theory

**DOI:** 10.1101/2025.06.19.660545

**Authors:** Dara Neumann, Qolamreza Razlighi, Yaakov Stern, Davangere P Devanand, Keith Jamison, Amy Kuceyeski, Ceren Tozlu

**Author notes:** **Corresponding Author:** Ceren Tozlu, PhD, **Mailing address:** 1300 York Avenue New York, NY 10065, **Email:**.

## Abstract

**Introduction:** Patients with mild cognitive impairment (MCI) have shown disruptions in both brain structure and function, often studied separately. However, understanding the relationship between brain structure and function can provide valuable insights into this early stage of cognitive decline for better treatment strategies to avoid its progression. Network Control Theory (NCT) is a multi-modal approach that captures the alterations in the brain’s energetic landscape by combining the brain’s functional activity and the structural connectome. Our study aims to explore the differences in the brain’s energetic landscape between people with MCI and healthy controls (HC).

**Methods:** Four hundred ninety-nine HC and 55 MCI patients were included. First, k-means was applied to functional MRI (fMRI) time series to identify commonly recurring brain activity states. Second, NCT was used to calculate the minimum energy required to transition between these brain activity states, otherwise known as transition energy (TE). The entropy of the fMRI time series as well as PET-derived amyloid beta (Aβ) and tau deposition were measured for each brain region. The TE and entropy were compared between MCI and HC at the network, regional, and global levels using linear models where age, sex, and intracranial volume were added as covariates. The association of TE and entropy with Aβ and tau deposition was investigated in MCI patients using linear models where age, sex, and intracranial volume were controlled.

**Results:** Commonly recurring brain activity states included those with high and low amplitude activity in visual (+/−), default mode (+/−), and dorsal attention (+/−) networks. Compared to HC, MCI patients required lower transition energy in the limbic network (adjusted *p* = 0.028). Decreased global entropy was observed in MCI patients compared to HC (*p* = 7.29e-7). There was a positive association between TE and entropy in the frontoparietal network (*p* = 7.03e-3). Increased global Aβ was associated with higher global entropy in MCI patients (ρ = 0.632, *p* = 0.041).

**Conclusion:** Lower TE in the limbic network in MCI patients may indicate either neurodegeneration-related neural loss and atrophy or a potential functional upregulation mechanism in this early stage of cognitive impairment. Future studies that include people with AD are needed to better characterize the changes in the energetic landscape in the later stages of cognitive impairment.

## Introduction

Mild cognitive impairment (MCI) is a clinical cognitive deficit with a prevalence ranging from 3% to as high as 42% in older adults in population-based studies (Tricco et al., 2012) and the prevalence increases with age. MCI patients have an increased risk of developing AD (Morris et al., 2001), therefore a diagnosis of MCI may represent a critical turning point in the trajectory of developing AD and be an opportunity for early intervention. Establishing neuroimaging signatures of MCI may allow more informed diagnosis, and, more importantly, an understanding of its underlying mechanisms that could pave the way for novel treatments.

Structural magnetic resonance imaging (MRI) has been largely used to identify neurodegeneration-related brain atrophy in MCI patients compared to controls (Frisoni et al., 2010). Previous studies have shown that multiple brain regions are impacted during the transition to the MCI phase, therefore identification of the changes at the network level would provide more sensitive biomarkers compared to single region-wise anatomical changes. Diffusion MRI (dMRI) and functional MRI (fMRI) are promising tools to identify the network-level changes at the microstructural and functional level even before the atrophy happens (Brier et al., 2014; Dai et al., 2019; Filippi et al., 2020; Gonuguntla et al., 2022; Hojjati et al., 2018). Additionally, the combination of dMRI and fMRI was also used to identify changes in brain structure and function coupling in different subnetworks, demonstrating the asymmetrical changes in the brain structure-function relationship in MCI (Sun et al., 2024).

Network Control Theory (NCT) is another approach that allows us to identify the relationship between brain structure and function. NCT is a multi-modal approach that measures the brain’s energetic landscape using the functional MRI time series and structural connectomes derived from diffusion MRI. Through NCT, we can measure the minimum energy required to transition between pairs of brain activation states, also known as transition energy (TE). Understanding the energetic landscape and its network-level locations serve as a critical marker for non-invasive therapies, such as transcranial magnetic stimulation (TMS), helping to identify the optimal stimulation sites to potentially delay or slow the progression of cognitive impairment. NCT has been used to identify changes in the brain’s energetic landscape due to disorders such as multiple sclerosis (Tozlu et al., 2023; Broeders et al., 2024), in subjects with heavy alcohol use (Singleton, Velidi, et al., 2023), in adolescents with a family history of substance use disorder (Schilling et al., 2024), as well as psychedelics’ effects on brain dynamics (Singleton et al., 2022; Singleton, Wang, et al., 2023).

However, to our knowledge, NCT has not been applied in the context of MCI or AD. Therefore, the primary goal of this study is to test for differences in the energy landscape in MCI patients compared to healthy controls (HC). We first identified the most commonly recurring brain activity states by clustering the fMRI time series. Second, we applied NCT approach using the fMRI time series and structural connectomes to estimate the minimum energy required to transition between each pair of dynamic brain states or to remain in the same state. Third, we compared the state-wise, regional, and global transition energy between MCI patients and HC. Additionally, we also computed the entropy of the regional fMRI time series and investigated the interrelationship between the entropy, transition energy, and positron emission tomography (PET) derived metrics of misfolded proteins (amyloid beta [Aβ] and tau deposition). Based on our previous work in disease/disorders, we hypothesized that greater energy would be required to transition between the brain states in MCI patients compared to controls. Understanding the altered energetic landscape of the brain in MCI patients may help to identify early signatures of MCI that can be used to identify individuals at risk of dementia or even to develop non-invasive therapies such as TMS to delay the progression to AD.

## Material and Methods

### Subjects

Four hundred ninety-nine healthy people (age: 66.7, sex: 59.9% F) and fifty-five people with MCI (age: 71.9, sex: 50.9% F) were included in our study. The participants were collected at Weill Cornell Medicine and Columbia University’s Irving Medical Center. Participants underwent 3T MRI, Aβ-PET (18F-Florbetaben for HC and 18F-Florbetapir for MCI), and harmonized 18F-MK6240 tau-PET scans within 12 months. All studies were approved by an ethical standards committee on human experimentation and written informed consent was obtained from all patients. Healthy controls underwent medical, neuropsychological, and MRI evaluations to confirm they had no neurological or psychiatric conditions, cognitive impairments, major medical illnesses, or any MRI contraindications. Participants were excluded if they had contraindications to MRI, had ever been diagnosed with or were currently on medication for a neurological or psychological disorder, or had functional MRI acquisition failed for that participant.

### Image Acquisition, Processing, and Structural Connectome Extraction

#### Structural and Functional MRI Acquisition

All structural MRI scans were conducted using 3.0 Tesla MRI scanners. Each participant’s imaging session began with a scout localizer to determine positioning and field of view, followed by a high-resolution magnetization-prepared rapid gradient-echo scan. MRI was performed with a TR/TE (time of repetition/time of echo) of 7/2.6 ms, a flip angle of 12°, a matrix size of 256 × 256, and 176 slices with a thickness of 1 mm. Resting State Functional fMRI (rs-fMRI) was performed with a TR/TE of 2000/23 ms, flip angle of 77°, matrix size of 128 × 128, voxel size of 1.5×1.5×3 mm, and 40 axial slices, each lasting 10 min. Rs-fMRI scans were acquired with participants’ eyes open.

#### Structural and Functional MRI Processing

All structural MRI scans were processed with FreeSurfer 7.1.0 (http://surfer.nmr.mgh.harvard.edu) for automated segmentation and cortical parcellation (e.g., including segmentation and creation of an average gray matter mask) (Fischl et al., 2002, 2004) to derive regions of interest (ROIs) in each subject’s native space using the Schaefer atlas (Schaefer et al., 2018). Intracranial volume (ICV) was calculated using 3T T1-weighted structural MRI and BrainWash, an automatic multi-atlas skull-stripping software package (Doshi et al., 2013).

Resting-state fMRI data are processed first by slice timing correction which is applied to the raw fMRI time series to account for the difference in the acquisition delay between slices. At the same time, motion parameters are estimated on raw fMRI scans using rigid-body registrations performed on all the volumes in reference to the first volume. Then, the estimated motion parameters are applied to the slice timing corrected fMRI time series to get the re-aligned fMRI data. To deal with involuntary head motion we consider two strategies. First, we perform scrubbing using frame-wise displacement > 0.5 mm and root mean squared difference > 0.3 as proposed and validated by Power et al., 2012. Second, we regress out the average time series of the deep white matter regions, the ventricles, and framewise displacement. Next, the temporal bandpass filtering is performed with a 0.01<f<0.1 Hz cut-off frequency. No spatial smoothing is applied. Finally, the fMRI data are transferred to the MNI space by performing a non-linear registration between the participant’s structural scan and MNI template using ANTs.

#### PET Acquisition

For tau-PET, all subjects were injected with 185 MBq (5 mCi) ± 20% (maximum volume 10 mL) of 18F-MK6240 before imaging that was administered as a slow single IV bolus at 60 s or less (6 s/mL max. Imaging was performed as six 5-min frames for a 30-min PET acquisition, 90–120 min post-injection). For Aβ-PET, subjects underwent 18F-Florbetaben or 18F-Florbetapir PET scans. This scan consisted of four 5-min frames over 20 min of acquisition, starting 50 min for the 18F-Florbetapir and 90 min for 18FFlorbetaben tracers after injection of 8.1 mCi ± 20% (300 MBq), which was administered as a slow single IV bolus at 60 s or less (6 s/mL max).

#### PET Processing

The same 200 cortical ROIs from the Schaefer atlas that were used for the fMRI inter-regional connectional connectivity metrics were utilized for calculating Aβ- and tau-PET regional uptakes. The Aβ and tau regional standardized uptake value ratio (SUVR) were calculated by normalization to cerebellum grey matter. As previously published, the fully automated in-house pipelines were utilized to process the Aβ and tau PET images (Brickman et al., 2015; Gu et al., 2015; Oh et al., 2015, 2016; Tahmi et al., 2019). Briefly, both Aβ- and tau-PET dynamic frames (six frames in tau-PET and four in Aβ-PET) were aligned to the first frame using rigid-body registration and averaged to generate a static PET image. Next, the structural T1 image in FreeSurfer space was registered to the same subject’s PET composite image using normalized mutual information and six degrees of freedom to obtain a rigid-body transformation matrix. Finally, the SUVR value for Aβ was converted to centiloid standard values due to different tracers in Aβ-PET data (Klunk et al., 2015).

#### Structural Connectomes

Diffusion MRI from the Human Connectome Project-Aging (HCP-A) (Bookheimer et al., 2019) was used to create the average structural connectivity network for this project as the age range was similar (the mean age of the HCP-A dataset is 60.37 ± 15.68 and the mean age of the dataset used in this study is 67.2). Diffusion MRI was processed by deterministic tractography with SIFT2 global streamline weighting and regional volume normalization. Structural connectomes were extracted using the Schaefer atlas, the same atlas used for the functional MRI. The structural connectome is a 200×200 matrix with a zero diagonal for each subject in the HCP-A dataset. We then averaged the structural connectomes from 716 subjects in the HCP-A dataset to create the average structural connectivity network which was used in the Network Control Theory approach to measure the transition energy.

### Network Control Theory

#### Brain State Identification

First, the fMRI time series were z-score normalized at the subject level using the overall average and standard deviation of the entire region x time matrix for each subject. Second, the normalized regional fMRI time series were concatenated for all subjects including HC and MCI patients, and then used in the k-means clustering to identify commonly recurring dynamic brain states (See Figure 1). We used both the elbow criterion and the gain in explained variance to identify the optimal number of clusters, which we identified to be six (see Supplementary Figure 1). Once the optimal number of clusters was identified, k-means clustering was run 10 times. Within each of the 10 runs, clustering was restarted 50 times with different random initializations to identify the final clusters having the best separation, that is, those that minimized Pearson’s correlation between pairs of cluster centroids. Adjusted mutual information (AMI) (Xuan Vinh Nguyen, 2024) was computed to assess the stability of the clustering results over the 10 repetitions; AMI was greater than 0.987 for all pairs of repetitions indicating robust cluster assignment (see Supplementary Figure 2).

**Figure 1:**
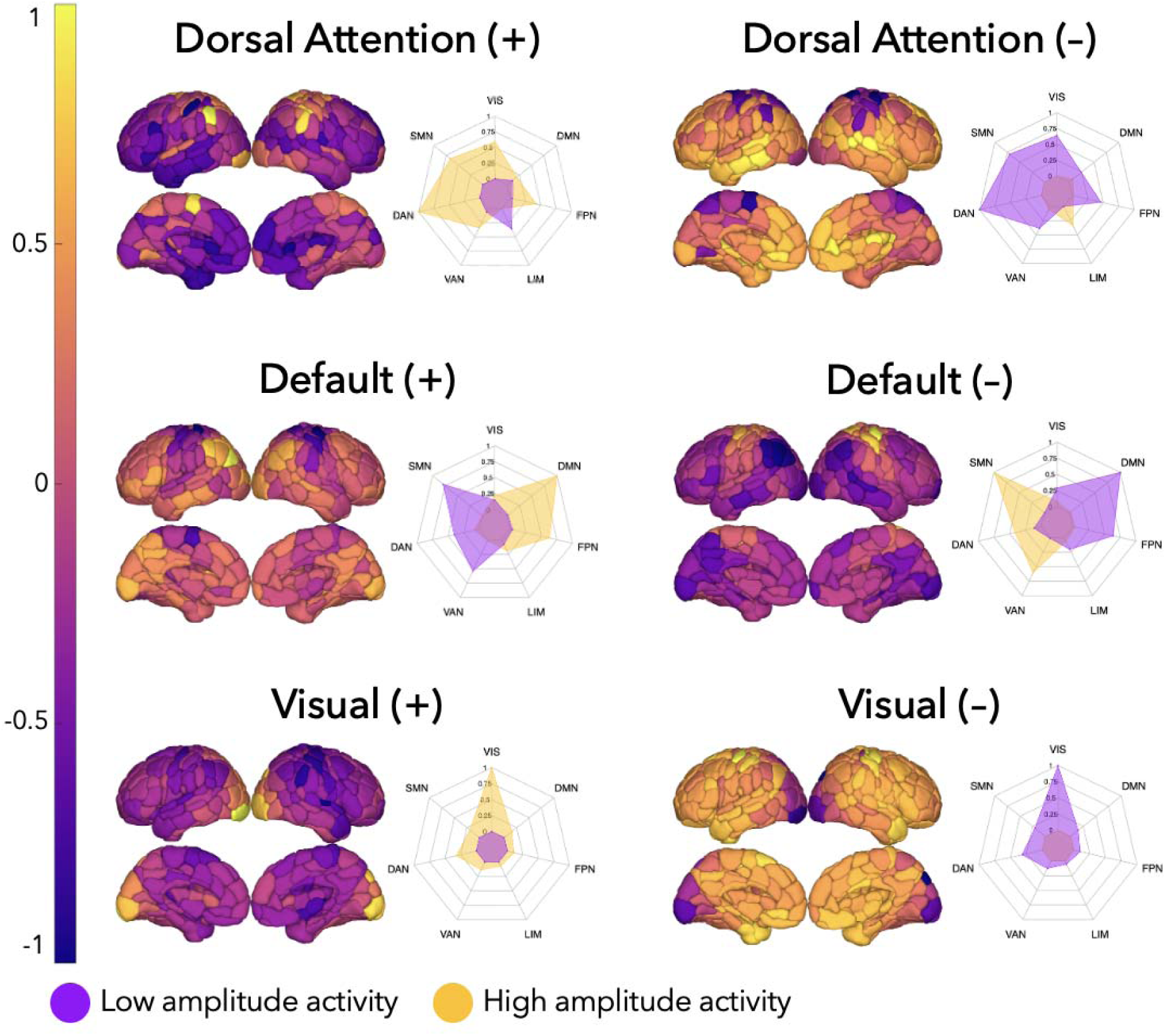
Brain plots and radial plots used to visualize the most recurring brain activity states. The radial plots show the mean of the positive and negative functional activity values over the Yeo functional network. States were named based on the network having the maximum magnitude in the radial plot. Visual (VIS), Dorsal Attention (DAN), Somatomotor (SMN), Salient Ventral Attention (VAN), Frontoparietal Control (FPN), Limbic (LIM), Default Mod (DMN) Network.

Each of the 200 regions was assigned to one of the seven Yeo functional networks (Yeo et al., 2011). The network-level contributions of the six brain states were characterized by calculating the cosine similarity of each state’s centroid (positive and negative parts were analyzed separately) and seven binarized vectors representing each of the networks. Because the mean signal from each scan’s regional time series was removed, positive values reflect signal intensity above the mean (high amplitude) and negative values reflect signal intensity below the mean (low amplitude). The brain states were named by identifying the network with the largest magnitude activation (either high+ or low– amplitude).

#### Transition Energy Calculations

As previously described (Cornblath et al., 2020; Singleton et al., 2022; Tozlu et al., 2023), NCT can be used to understand how the structural connectome constrains dynamic brain state changes. Specifically, it can be used to compute the minimum energy (*E_m_*) required to transition between pairs of brain states or to remain in the same state. To begin, we employed a time-invariant model:

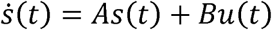

where *A* is the structural connectivity matrix with a dimension of 200 × 200, *s*(*t*) is a vector of length 200 containing the regional activity at time *t*, and *u*(*t*) is the external input to the system. In our case, *B* is an identity matrix with a dimension of 200 × 200 that indicates each region has uniform control over the system.

When computing the transition energy (*E_m_*) from the initial activity *s*_0_ to final activity *s_f_* over some time *T*, we use an invertible controllability Gramian *W* to control the structural connectivity network *A* from the set of network nodes *B*. The invertible controllability Gramian *W* is defined as:

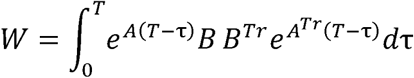

where *T_r_* denotes the matrix transpose. Optimal T was identified by grid searching between 0.001 and 10 and identifying the value that maximized the absolute value of Spearman’s correlation between the entries in the transition probability matrix and the entries in the transition energy matrix. The correlation between transition probability and energy is expected to be negative since the brain prefers trajectories through state space requiring minimal input energy given structural constraints, therefore the brain tends to change less between states if the minimum required energy is greater. Here, we identified an optimal T of 0.501, which yielded a correlation coefficient of −0.763, i.e. absolute value of the correlation coefficient: 0.763 (*p* = 5.22e-7) (see Supplementary Figure 3). Additionally, we replicated our results with T = 1 as previously used in another study (Singleton et al., 2024) (See Supplementary Figures 7 and 8).

After computing the invertible controllability Gramian W and optimizing T, the transition energy (*E_m_*) is then calculated as the quadratic product between the inverted controllability Gramian W and the difference between *e^AT^s*_0_, which represents the activation at time T assuming brain activity spreads in a network diffusion manner over the SC from the initial state *s*_0_, and the final state *s_T_*: *E_m_* = (*e^At^s_0_* - *s_T_*)*^Tr^ W*-1(*e^AT^s*_0_ - *s_T_*).

Since there were 6 brain states, the transition energy matrix is of size 6 × 6 where off-diagonal entries contain the energy required to transition between each pair of unique states *s*_0_ and *s_T_*, where *s*_0_ ≠ *s_T_*, and diagonal entries contain persistence energy required to stay in the same state (*s*_0_ – *s_T_*).

### Entropy of fMRI Brain Activity

To investigate how TE was related to the amount of information contained in the fMRI brain activity signal, regional entropy was calculated directly on the fMRI time series. Denoting the regional fMRI time series in a region by *s*(*t*) - (*s*(*1*),…, *s*(*n*)) where *n* is the number of the fMRI time series acquisition. We define two template vectors of length m such as: *s_m,i_* = (*s*(*i*), *s*(*i* +1),…,*s*(*i* + *m*-1)) and *s_m,j_* = (*s*(*j*), *s*(*j* + 1),…, *s*(*j* + *m* - 1)). Entropy was calculated as the negative logarithm of the probability that if two template vectors of length m have a distance less than *r* (i.e., *d*[*s_m,i_,s_m,j_*] < *r*) then two template vectors of length *m* + 1 also have a distance less than r (i.e., *d*[*s_m +1,i_,s_m+1,j_*] < *r*) such as

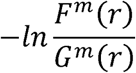

where *F_m_*(*r*) is the number of template vector pairs having *d*[*s_m +1,i_,s_m+1,j_*] < *r*, *G_m_*(*r*) is the number of template vector pairs having *d*[*s_m,i_,s_m,j_*] < *r* and where *d* was the Euclidean distance (Richman & Moorman, 2000). In our study, *m* = 3, as suggested in three previous in people with multiple sclerosis (Tozlu et al., 2023; Zhou et al., 2016). The parameter *r* was studies that computed entropy from resting state fMRI data in controls (Z. Wang et al., 2014) and defined as 0.2*sd*(*s*) where *sd*(*s*) is the standard deviation of the time series (s), as recommended by previous work (Tomčala, 2020). An individual’s global entropy was calculated as the average of their regional entropies.

### Statistical Analysis

The comparison of neuroimaging biomarkers (i.e., transition energy, entropy, tau, and Aβ metrics) was performed using linear regression models where the output was the neuroimaging variable, and the inputs were the groups (HC vs. MCI), age, sex, and ICV. The relationship between each pair of neuroimaging biomarkers (TE vs. PET metrics and entropy vs. PET metrics) was assessed using linear regression models where age, sex, and intracranial volume were added as covariates. The p-values obtained with multiple comparisons for the region, network, and state-wise analysis were corrected using the Benjamini-Hochberg method. The results were considered significant when the corrected *p*-value < 0.05. All statistical analyses and graphs were performed using R (https://www.r-project.org) version 4.2.2 and MATLAB (https://www.mathworks.com/) version R2023b.

## Results

### Subject Characteristics

Our cohort consisted of 554 subjects including 55 MCI and 499 HC (See Table 1). While sex and ICV were not significantly different between HC and MCI (*p* = 0.25 and *p* = 0.10, respectively), the MCI group was older compared to HC (*p* = 3.5e-7).

**Table 1:** The age, sex, and ICV information for HC and MCI patients. The mean and standard deviation of age and ICV are reported, while count and percent were reported for sex. The p-values were computed using the Wilcoxon rank sum test to compare age and ICV, and the Chi-Squared test was used to compare the sex between HC and MCI. The * indicates significant *p*-values, i.e. *p*-value < 0.05.

|  | All Participants<br>(n = 554) | HC<br>(n = 499) | MCI<br>(n = 55) | *p*-value<br>(HC vs. MCI) |
| --- | --- | --- | --- | --- |
| Age | 67.2 ± 5.99 | 66.7 ± 5.53 | 71.9 ± 7.77 | 3.249e-7* |
| Sex (F (%)) | 327 (59.0%) | 299 (59.9%) | 28 (50.9%) | 0.2521 |
| ICV (L) | 1.45 ± 1.52 | 1.44 ± 1.51 | 1.49 ± 1.60 | 0.1013 |

### Brain States

Figure 1 shows the most recurring brain states obtained by clustering the fMRI time series with the k-means approach (k = 6). The brain states consist of three pairs of anticorrelated states with high/low-amplitude activity: dorsal attention (DAN+/–), default mode (DMN+/–), and visual (VIS+/–) networks.

### Comparison of Brain Energetics

Figure 2 shows the global, state-wise, network-wise, and regional TE in MCI and HC, as well a the beta estimates obtained from the linear model comparing these metrics between the MCI and HC groups. Global TE was not significantly higher in HC compared to MCI (*p* = 0.796), however TE in the limbic network was significantly higher in HC compared to MCI (adjusted *p* = 0.028). Additionally, the TE was significantly higher in 4 brain regions in HC compared to MCI, while it was higher in 26 regions in MCI compared to HC after multiple comparison correction. Finally, we replicated the TE analyses with T = 1. The results obtained with T = 0.501 were similar to those with T = 1 (See Supplementary Figure 5).

**Figure 2:**
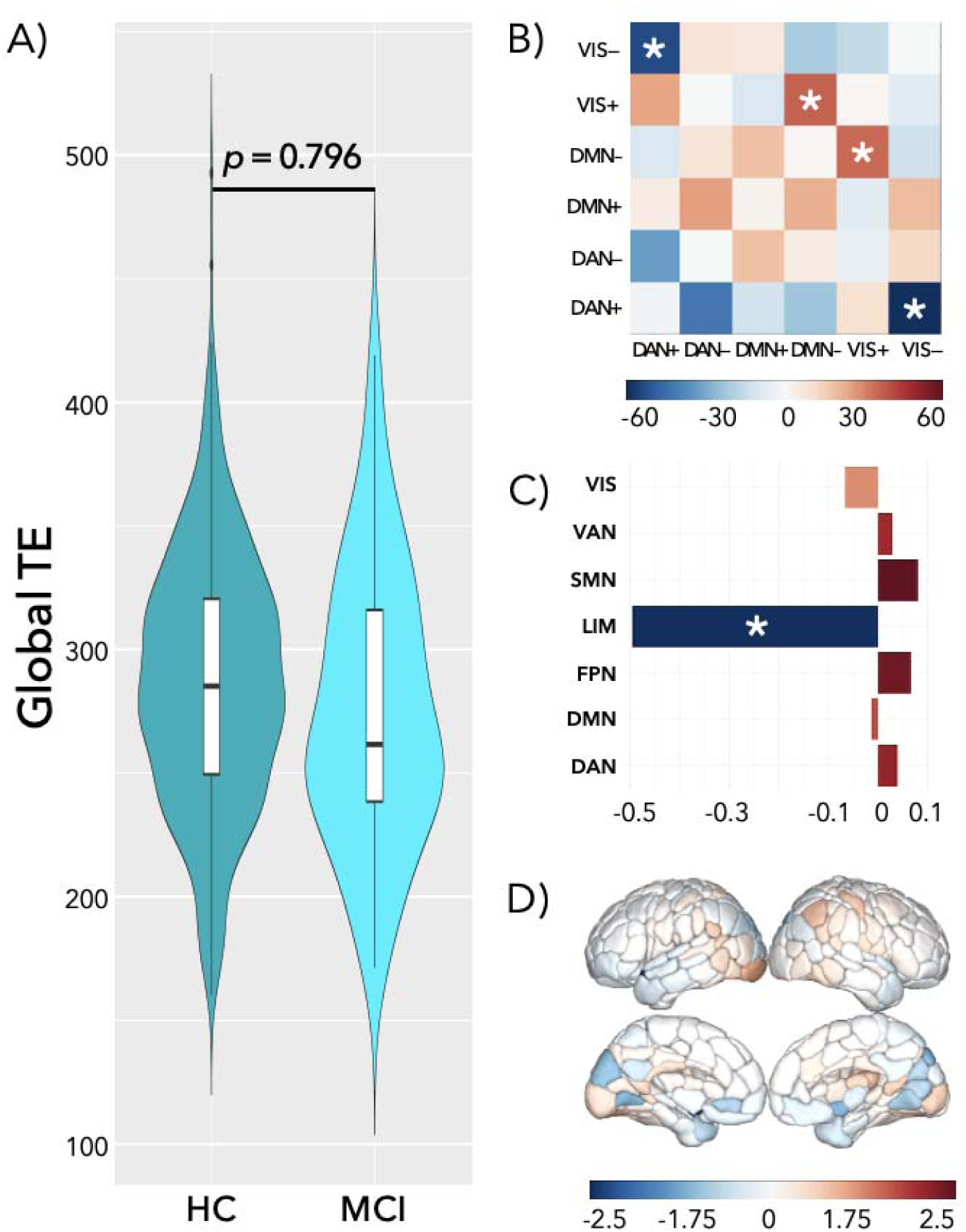
(A) Distribution of global transition energy in HC and MCI. The p-value (*p* = 0.796) was computed using the linear model where the output was the global TE and the inputs were the group assignment (MCI vs. HC), age, sex, and ICV. (B) The beta estimates from the linear model adjusted for age, sex, and ICV were used to represent the amplitude and the direction of the difference in the state-wise TE between HC and MCI groups. State pairs with significantly different TE before correction are marked with an asterisk. None of the state pairs were significant after correction. (C) The beta estimates from the linear model adjusted for age, sex, and ICV were used to represent the amplitude and the direction of the difference in the network-wise TE between HC and MCI groups. The limbic network (marked with *) was significant after multiple comparison *p*-value corrections. (D) The beta estimates from the linear model adjusted for age, sex, and ICV are used to represent the amplitude and the direction of the difference in the regional TE between HC and MCI groups. In figures B-D, positive values represent higher TE in MCI compared to HC, while negative values represent higher TE in HC compared to the MCI group.

### Comparison of entropy between MCI vs. HC

Figure 3 shows the global entropy calculated in MCI and HC separately as well as the beta estimates from the linear model comparing the regional entropy between MCI and HC groups. The global entropy was significantly higher in HC compared to MCI (*p* = 7.29e-7). One hundred ninety-nine out of 200 regions had significantly higher entropy in HC compared to MCI after the multiple comparison *p*-value correction.

**Figure 3:**
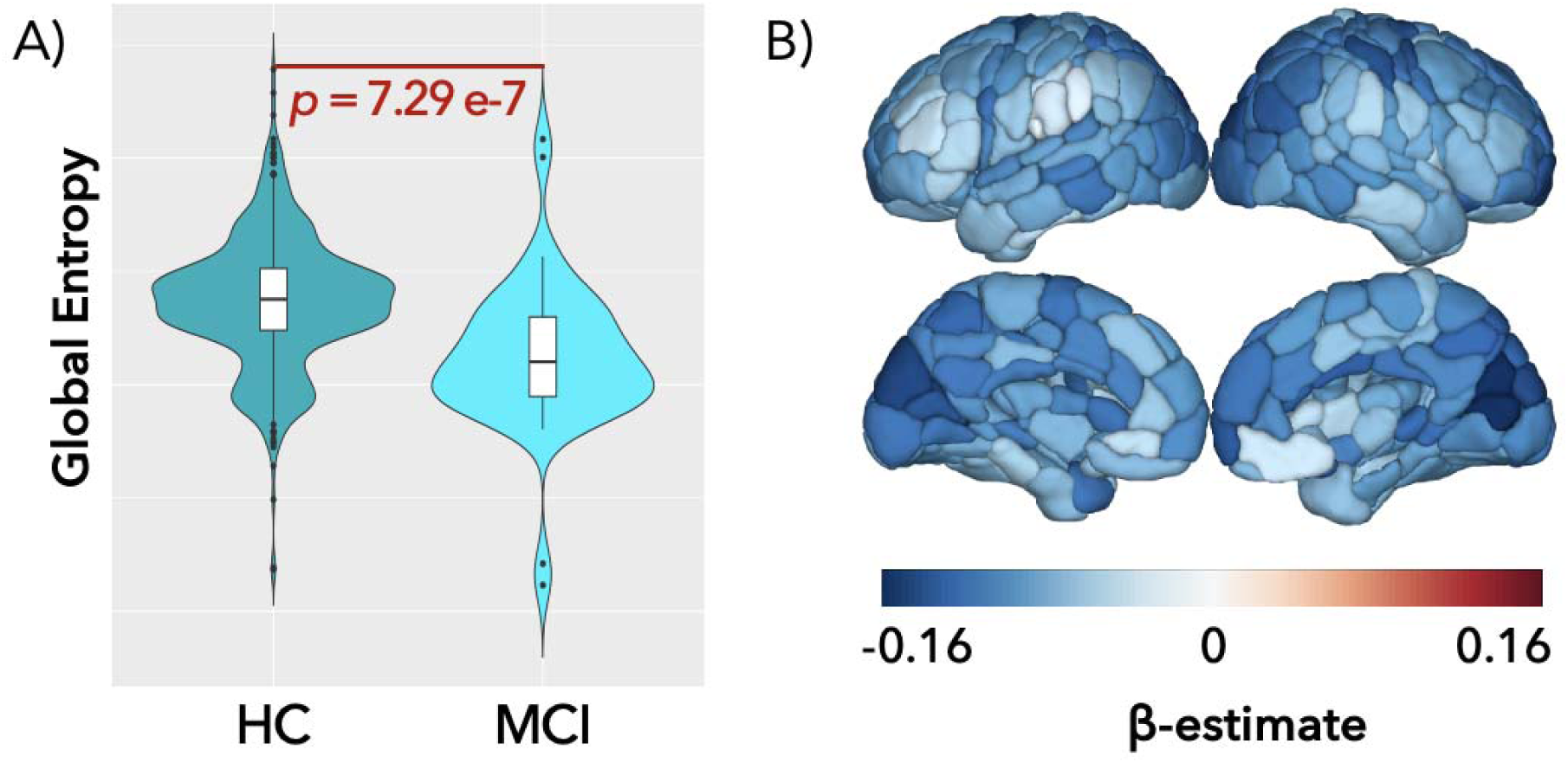
(A) Distribution of global entropy in HC and MCI. (B) The beta estimates from the linear model adjusted for age, sex, and ICV were used to represent the amplitude and the direction of the difference in the regional entropy between HC and MCI groups. Positive values represent higher regional entropy in MCI compared to HC, while negative values represent higher regional entropy in HC compared to the MCI group.

**Figure 4:**
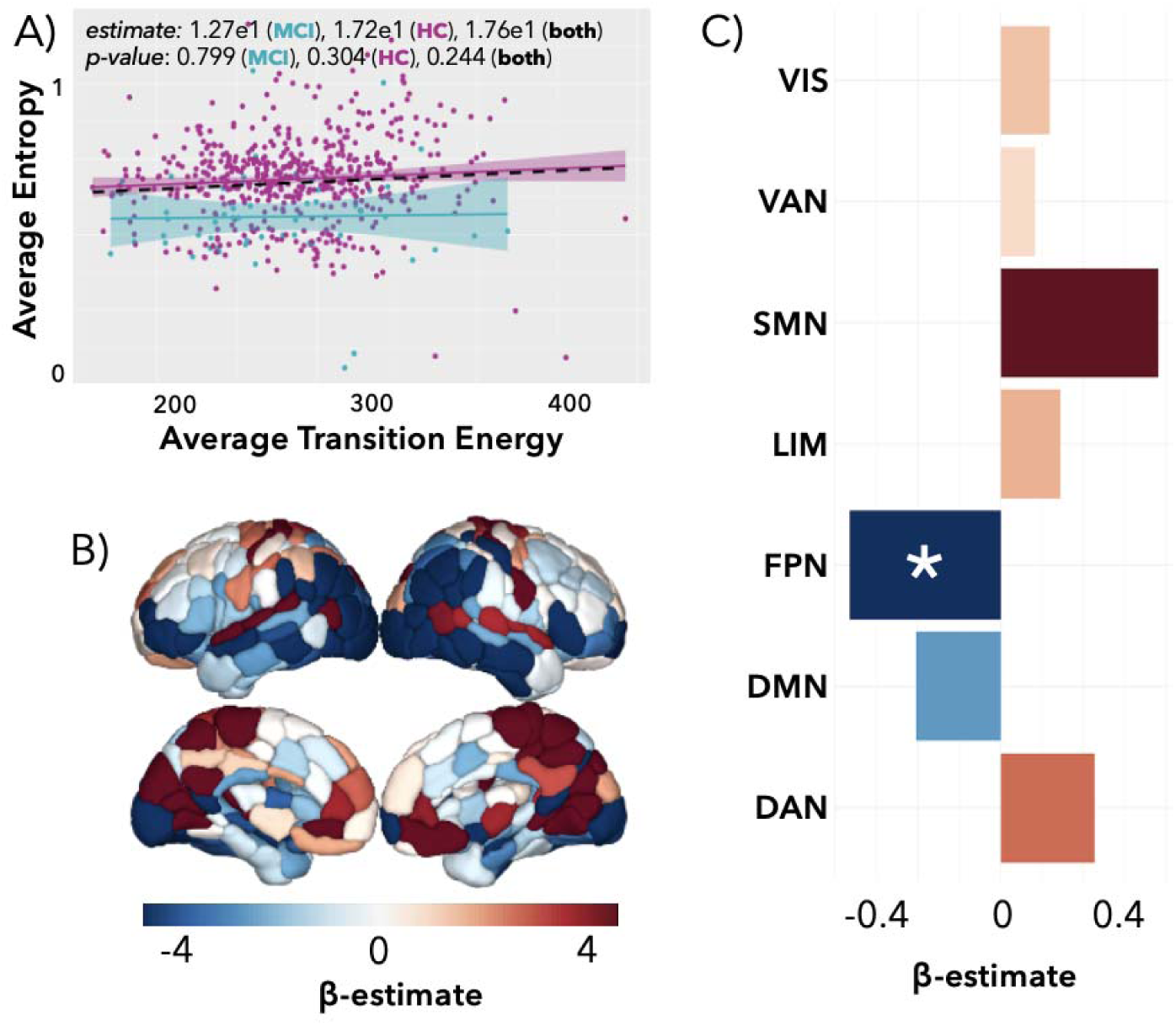
(A) The scatterplot shows the association between average entropy and average transition energy in all subjects, and in HC and MCI groups separately. (B) The beta estimates from the linear model adjusted for age, sex, and ICV were used to represent the amplitude and direction of association between global entropy and global transition energy. (C) The beta estimates from the linear model adjusted for age, sex, and ICV were used to represent the amplitude and the direction of the association between network-wise TE and network-wise entropy. The frontoparietal control network (marked with an asterisk) was significant after multiple comparison *p*-value corrections.

### Association between transition energy and entropy

While there was no association between global TE vs. entropy, there were significant associations at the region level. 29 regions showed a significant association between regional TE and entropy. Of these 29 regions, 13 showed a significant positive association between TE and entropy and 16 showed a negative association. At the network level, TE was negatively associated with entropy in the frontoparietal network (BH corrected *p*-value = 7.03e-3).

### Comparison of transition energy, entropy, and PET metrics

As expected, Aβ plaque and tau protein levels were higher in most regions in the MCI group compared to controls (See Supplementary Figure 5). Figure 5 shows the association of PET metrics with TE and entropy at the global level in MCI patients (See Supplementary Figure 5 for both MCI and HC). Higher global Aβ plaque level was associated with increased global entropy in MCI patients (*p* = 0.041). The TE obtained with T = 1 showed similar results (See Supplementary Figure 8). We also explored the associations of regional PET metrics with regional TE and entropy; however, these associations did not remain significant after *p*-valu correction for multiple comparisons (see Supplementary Figure 6).

**Figure 5:**
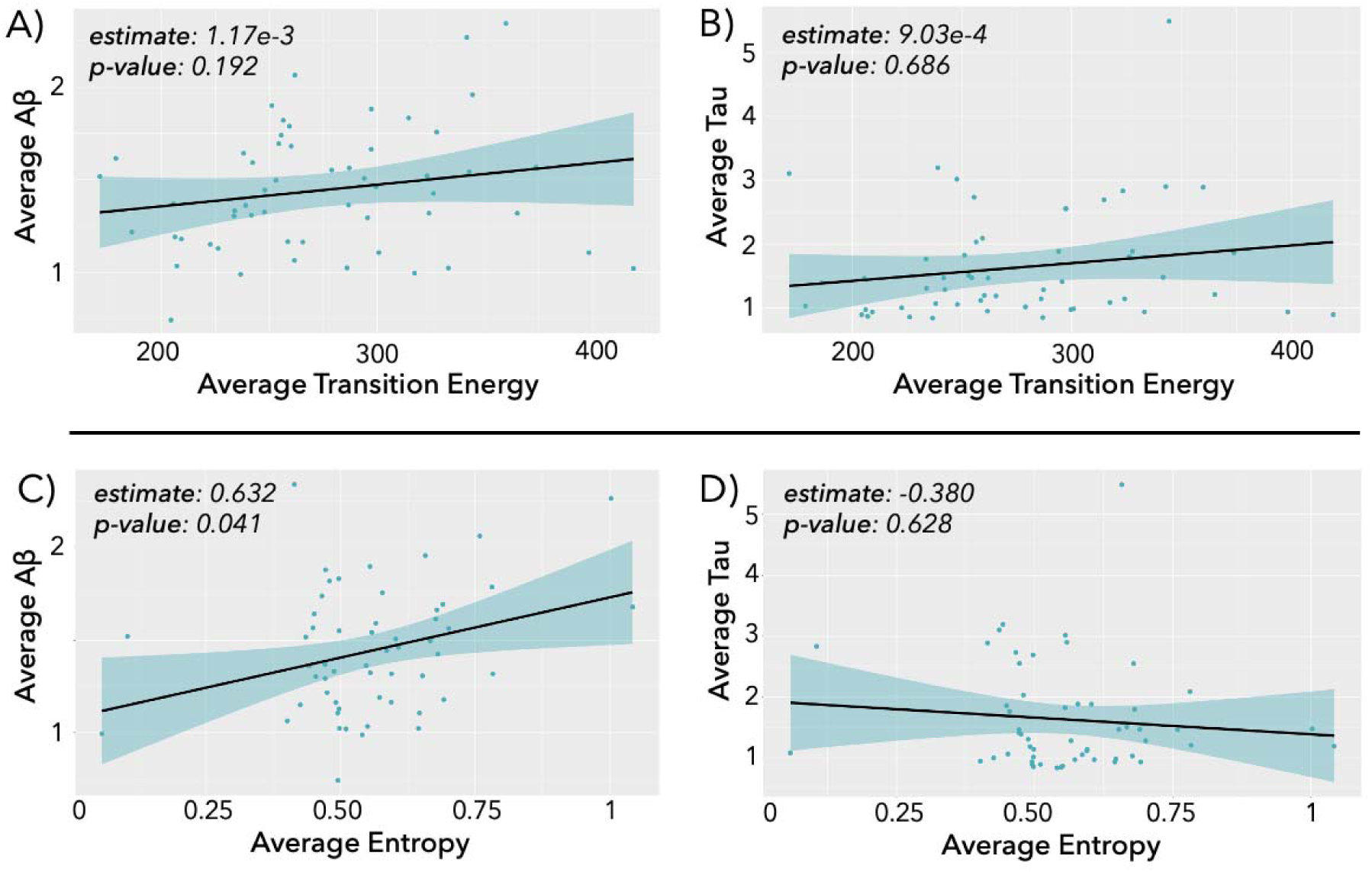
The scatter plots represent the association between (A) global TE and global tau protein levels, (B) global TE and global Aβ plaque levels, (C) global entropy and global tau protein levels, and (D) global entropy and global Aβ plaque levels. The data shown in this figure only represents MCI patients. Aβ plaque and tau protein levels are given in units of standard uptake value ratio (SUVR). The β and the *p*-value from the linear model adjusted for age, sex, and ICV are presented on the scatter plots.

## Discussion

The primary goal of our study was to identify the differences in the energetic landscape and entropy in MCI patients compared to HC. Additionally, we explored the association between these metrics and PET metrics such as Aβ and tau in MCI patients. Our key findings were 1) TE in the limbic network was significantly lower in MCI compared to HC, 2) decreased global and regional entropy was observed in MCI patients compared to HC, 3) increased global Aβ wa associated with greater global entropy in MCI patients. Taken together, the reduced limbic network energetic cost observed in MCI compared to HC may reflect underlying neural loss and atrophy due to neurodegeneration, leading to decreased metabolic demand at the network level. Alternatively, it may also suggest a functional upregulation mechanism, whereby the brain increases functional activity to compensate for structural damage, enabling more efficient transitions between brain states with lower energetic requirements in MCI patients, who are in the early stage of the cognitive impairment spectrum.

### A potential compensation mechanism in MCI patients

Lower energetic need to transition between states in MCI patients might be due to either (1) synaptic dysfunction and/or neural loss which may decrease the functional activity, therefore energetic demand, or (2) increased functional upregulation to compensate the pathological accumulation such as Aβ and tau, that may result in easier shifts between dynamic brain states with lower energetic demand. Lower metabolic energy as measured with the Fluorodeoxyglucose-PET (FDG-PET) has been observed in MCI patients compared to HC, particularly in the limbic and temporal lobes (Sanabria-Diaz et al., 2013). It has been speculated that lower energetic demand measured with FDG-PET is associated with synaptic dysfunction, a hallmark of early-stage neurodegeneration. Moreover, FDG-PET metrics have been shown to correlate with TE measured by NCT (He et al., 2022). Altogether, our findings of reduced TE in MCI patients may reflect early synaptic dysfunction, potentially marking the initial stages of neurodegenerative processes.

The compensation mechanism has been also observed in many previous studies that used fMRI. Another study showed that people with less severe MCI showed increased fMRI activity during associative recognition only, while people with more severe MCI showed increased fMRI activity during item recognition only, demonstrating a potential compensatory mechanism (Clément & Belleville, 2012). Our study builds on these findings by showing that increased functional upregulation may result in easier transitions between the brain states with decreased overall energetic demand in MCI patients compared to controls.

### Decreased energetic demand in the specific networks

The TE required in the limbic network was lower in MCI patients compared to HC. Previous studies have shown increased global network synchronization in MCI (Buldú et al., 2011), as well as a specific increased activation in the hippocampus and medial temporal lobe, which are a part of the limbic system, in MCI patients compared to HC and people with AD (Dickerson et al., 2004, 2005). Increased functional activity in those regions might result in decreased energetic demand within the limbic network.

### Decreased entropy in MCI patients

The entropy of the functional brain activity signals is being largely used to identify the abnormalities in signal complexity and its relationship with other neuroimaging biomarkers in people with neurological disorders. Our current study showed that global or regional entropy is lower in MCI compared to HC. Similar to these results, our group has also shown that lower entropy of the fMRI metrics is associated with larger lesion volume in people with MS (Tozlu et al., 2023). Previous studies on MCI and AD also found lower entropy of the fMRI metrics in MCI patients and AD compared to HC. A study that used permutation entropy, rather than sample entropy used in this study, found decreased entropy in the left cuneus in the MCI group compared to HC (Wang et al., 2017). Another study that used multi-scale entropy approach found decreased entropy in various regions (Niu et al., 2018), confirming our results that showed 199 out of 200 regions had lower entropy in MCI patients compared to HC.

### Association between entropy and A**β** levels in MCI patients

Our results showed that higher global Aβ accumulation is associated with increased entropy in patients with MCI. Previous studies have also shown that increased Aβ accumulation is associated with increased functional connectivity in the default mode, frontoparietal and motor networks in a longitudinal study that included MCI patients (Moffat et al., 2022). Another study also reported that elevated Aβ levels may differentially impact excitatory and inhibitory synapses, leading to abnormal network activity patterns and a potential increase in functional activity (Palop & Mucke, 2010). Our group also recently showed using a longitudinal dataset that increase in Aβ levels are associated with elevated functional connectivity of limbic with the default mode and control networks (Hojjati et al., 2025). The observed positive association between the Aβ level and entropy in our study could be attributed to disrupted functional activity with higher unpredictability and variability as a response to Aβ accumulation.

### Limitations

One of the limitations of our study was that we did not have individual structural connectomes available. Therefore, we relied on the averaged structural connectomes from the HCP-A dataset when computing the TE, as the age range of the HCP-A dataset is similar to the cohort we used in this study. Our experience applying NCT across various datasets demonstrated that the TE is similar when computed using individual versus averaged structural connectomes, therefore we expect that the results would be similar if the individual structural connectomes were used. Another limitation of our study was that our data only included MCI patients. Future research incorporating people with both early and late AD is necessary to better understand the full spectrum of the changing energetic landscape from HC to AD. Finally, our study included only cross-sectional data, limiting our ability to map the longitudinal association between energetic landscape and cognitive impairment. A future longitudinal study is needed to better understand how the brain’s energetic dynamics evolve over time and how it relates to the cognitive decline seen in MCI patients, who are prone to developing AD.

## Conclusion

In summary, our study highlights significant alterations in the energetic landscape and entropy in MCI patients compared to HC, providing novel insights into the brain structure-function relationship underlying early cognitive impairment. These findings have important implications for understanding the early disruptions in the brain due to MCI and may inform future non-invasive therapeutic interventions such as TMS, aimed at modulating network activity and alleviating clinical symptoms, delaying the progression of the disease, and improving the quality of life for MCI patients.

## Acknowledgements

D.N. performed the statistical analysis, created the figures, and wrote the article.

Q.R. collected the data and edited the article.

Y.S. collected the data and edited the article.

D.D. collected the data and edited the article.

K.J. generated the structural connectomes from the HCP-A dataset.

A.K. designed and supervised the study and edited the article.

C.T. designed and supervised the study and edited the article.

## Data/code availability statement

The deidentified data that support the findings of this study are available upon reasonable request from the corresponding author. Please see the paper from Cornblath et al., 2020 for the codes that were used for the clustering and network control theory approach. We added our code that generated the results and figures of this study to https://github.com/dn3umann/nct-mci.

## Ethics statement

All studies were approved by an ethical standards committee of Weill Cornell Medicine and Columbia University on human experimentation, and written informed consent was obtained from all patients.

