## Supplementary Information for "Disrupted Energy Landscape in Individuals with Mild Cognitive Impairment: Insights from Network Control Theory"

**Supplementary Material**

Supplementary Figure 1 shows the variance explained and gain in variance explained results when k-means clustering was applied to cluster the fMRI time series with different number of clusters ranging between 2 and 10. For each number of clusters, variance explained was calculated using the centroids of the time series data for the respective number of clusters. The best number of clusters was determined to be k = 6 because increasing k beyond 6 resulted in a less than 1% increase in variance explained.

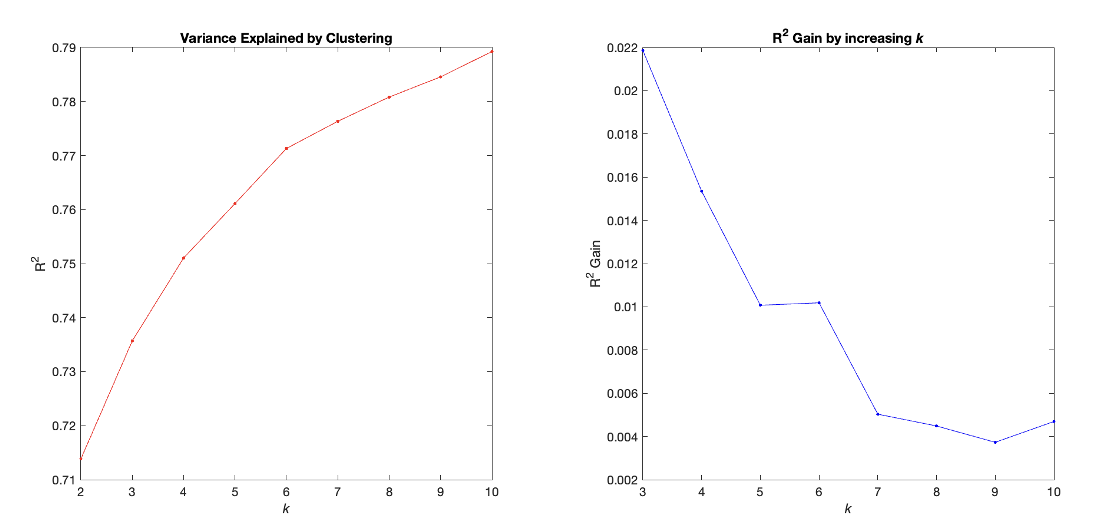

**Supplementary Figure 1**: **Choosing the best number of clusters**. Explained variance and gain in explained variance for values of k ranging between 2 and 10 clusters.

Supplementary Figure 2 shows the results of Adjusted Mutual Information (AMI) which informs about the stability of the clustering between 10 independently generated partitions of our data when 6 clusters were used. Our results show that the clusters were similar to each other with an AMI score > 0.98.

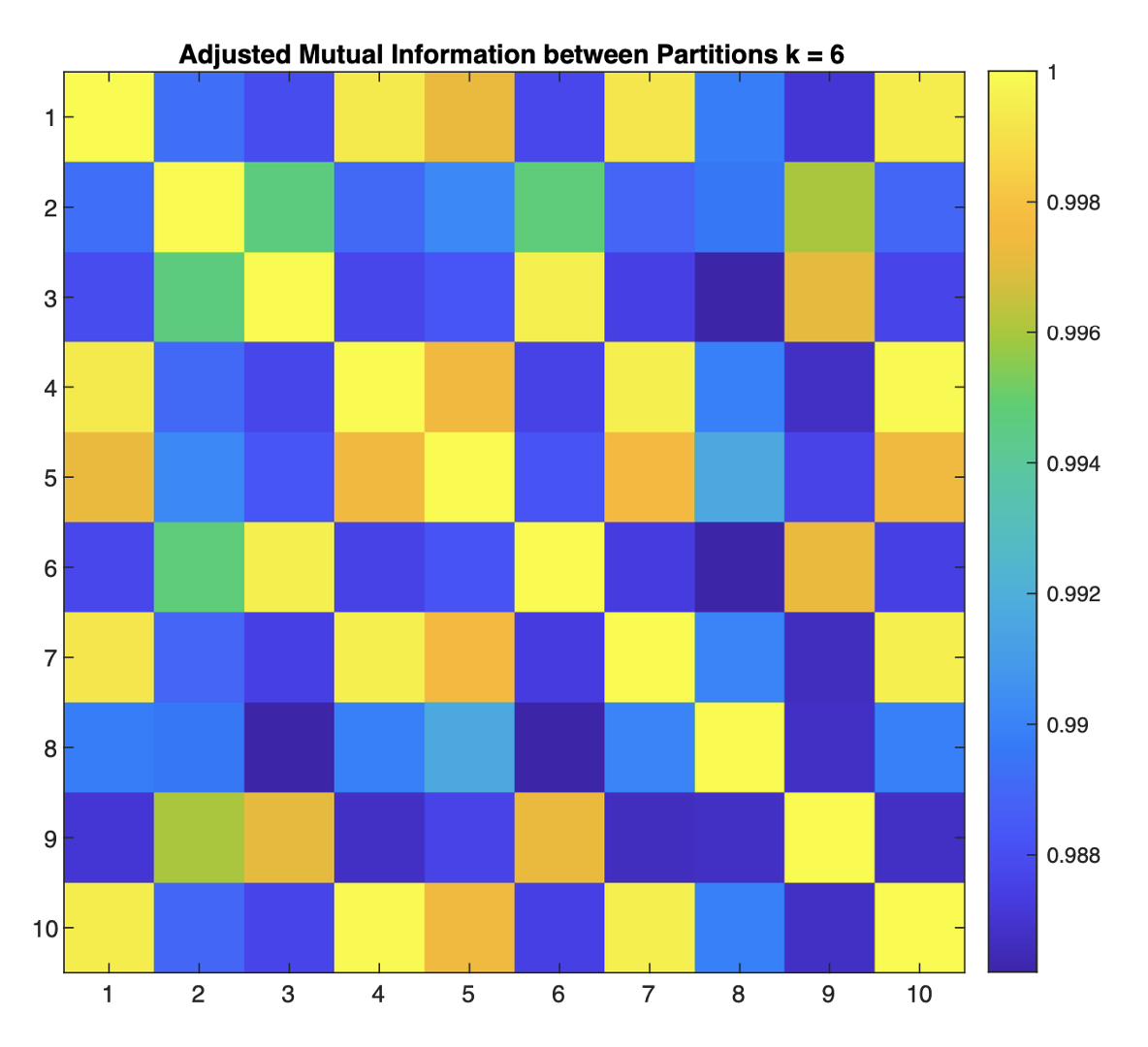

**Supplementary Figure 2: Assessment of clustering stability**. Adjusted mutual information (AMI) shared between 10 independently generated partitions of our data for k = 6. Values ranged between 0 and 1, where 1 indicates identical partitions.

Supplementary Figure 3 shows the amplitude of the Spearman correlation between the transition energy and transition probability using different values of T, ranging from 0.001 to 10 with steps of 0.5. The best value for T was T = 0.501 with an amplitude of correlation of 0.763 and a p-value of 5.22e-7. Therefore, T = 0.501 was used to compute minimum transition energy.

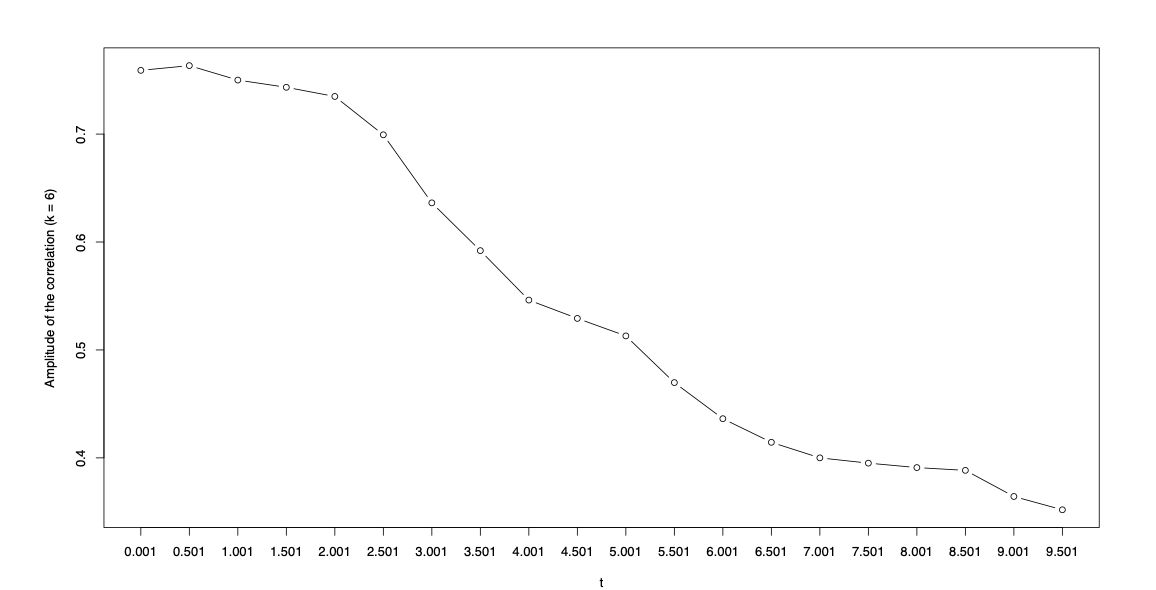

**Supplementary Figure 3: Optimization of the T parameter**. Correlation between transition energy and transition probability when varying the parameter T. The highest correlation was obtained for T = 0.501.

Supplementary Figure 4 shows the average regional and global Aβ and tau levels in MCI and HC groups separately as well as the β values comparing the regional plaques between MCI and HC. The statistically significant results were reported when the linear model adjusted for age, sex, and ICV yielded a *p*-value less than 0.05. 188 out of 200 regions had significantly different Aβ between HC and MCI (*p* < 0.05). 185 of these 188 regions had significantly higher Aβ in MCI than in HC. The global plaque was higher in MCI compared to HC. 199 out of 200 regions had significantly different tau levels between HC and MCI (*p* < 0.05). All 199 of these regions had significantly higher tau levels in MCI than in HC. The global tau level was higher in MCI compared to HC using linear model with age, sex, and ICV as covariates.

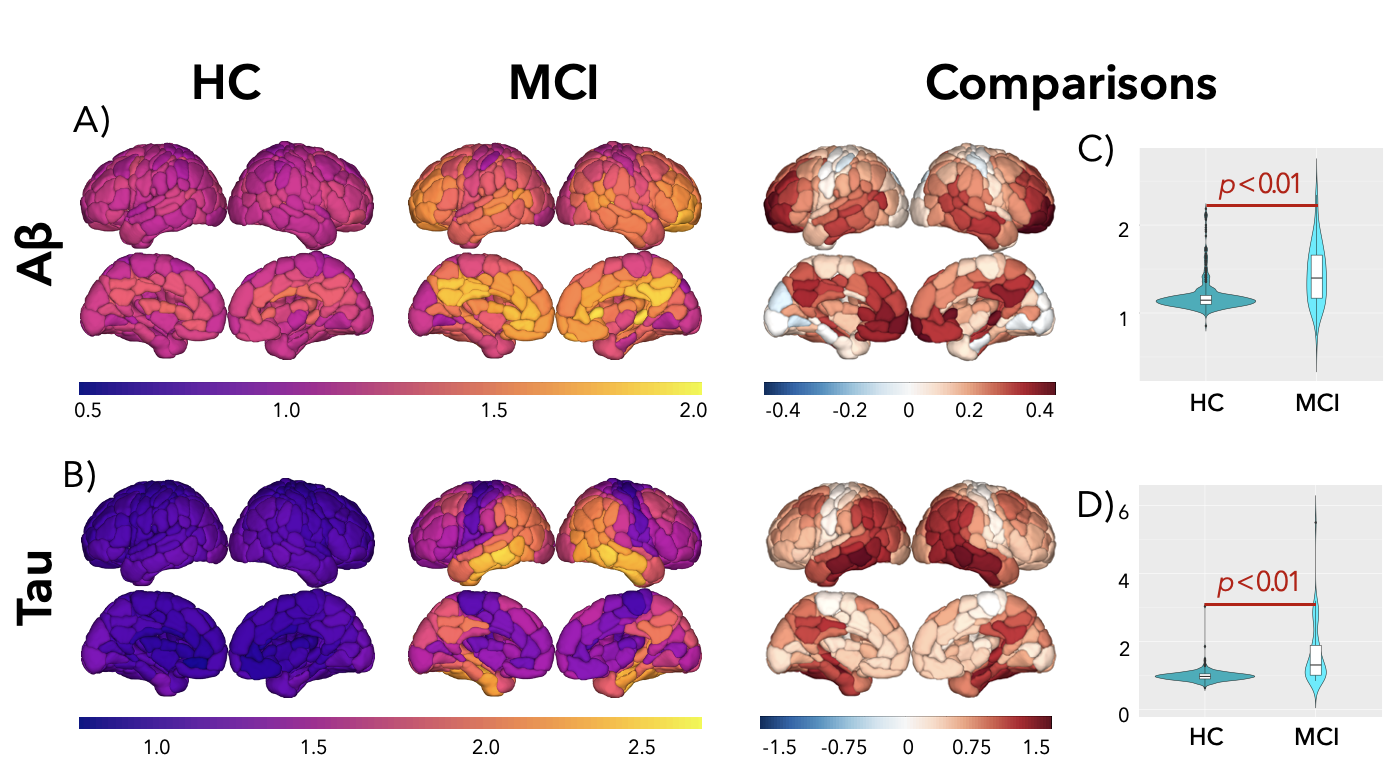

**Supplementary Figure 4:** (A) The average regional Aβ levels in HC and MCI groups separately, then the statistics from the linear model adjusted for age, sex, and ICV were used to represent the amplitude and the direction of the difference in the regional Aβ levels between HC and MCI groups. (B) The average regional tau levels in HC and MCI groups separately, then the statistics from the linear model adjusted for age, sex, and ICV were used to represent the amplitude and the direction of the difference in the regional tau levels between HC and MCI groups. The positive values represent higher regional Aβ or tau levels in MCI compared to HC, while negative values represent higher regional Aβ or tau levels in HC compared to the MCI group. Aβ and tau levels are given in units of standard uptake value ratio (SUVR). (C) Distribution of global Aβ levels in HC and MCI. (D) Distribution of global tau levels in HC and MCI. The *p*-values were computed using the linear model adjusted for age, sex, and ICV.

The Supplementary Figure 5 is the replicated version of Figure 5 by plotting the same metrics using both HC and MCI patients.

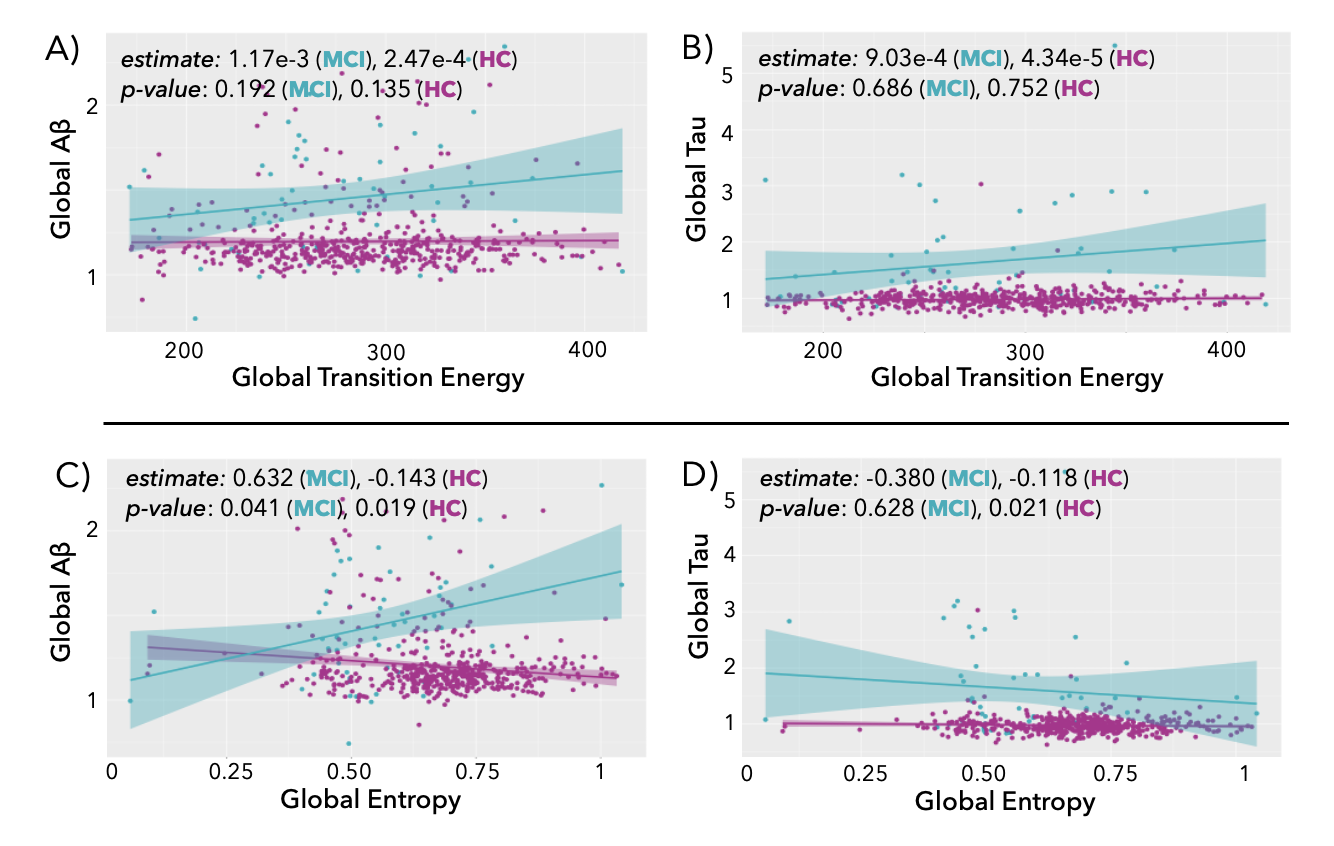

**Supplementary Figure 5**: The scatter plots represent the association between (A) global TE and global entropy, (B) global and global tau protein levels, (C) global TE calculated with T = 0.501 and global Aβ plaque levels, (D) global entropy and global tau protein levels, and (E) global entropy and global Aβ plaque levels. The data shown in this figure represents both HC and MCI patients. Aβ plaque and tau protein levels are given in units of standard uptake value ratio (SUVR). The β and the *p*-value from the linear model adjusted for age, sex, and ICV are presented on the scatter plots.

Supplementary Figure 6 shows the Spearman correlation between regional TE, entropy, Aβ, and tau in MCI patients. We only focused on MCI patients in this analysis as we were interested in the association of the MCI related changes in the PET with TE and entropy. There was a significant positive correlation between entropy and Aβ in two regions, one in the left orbitofrontal cortex region and the other in the right visual region, in which increased Aβ was associated with higher entropy. After multiple comparison p-value correction, there was no significant correlation between regional entropy with tau. Additionally, there was no correlation between regional TE and both PET metrics.

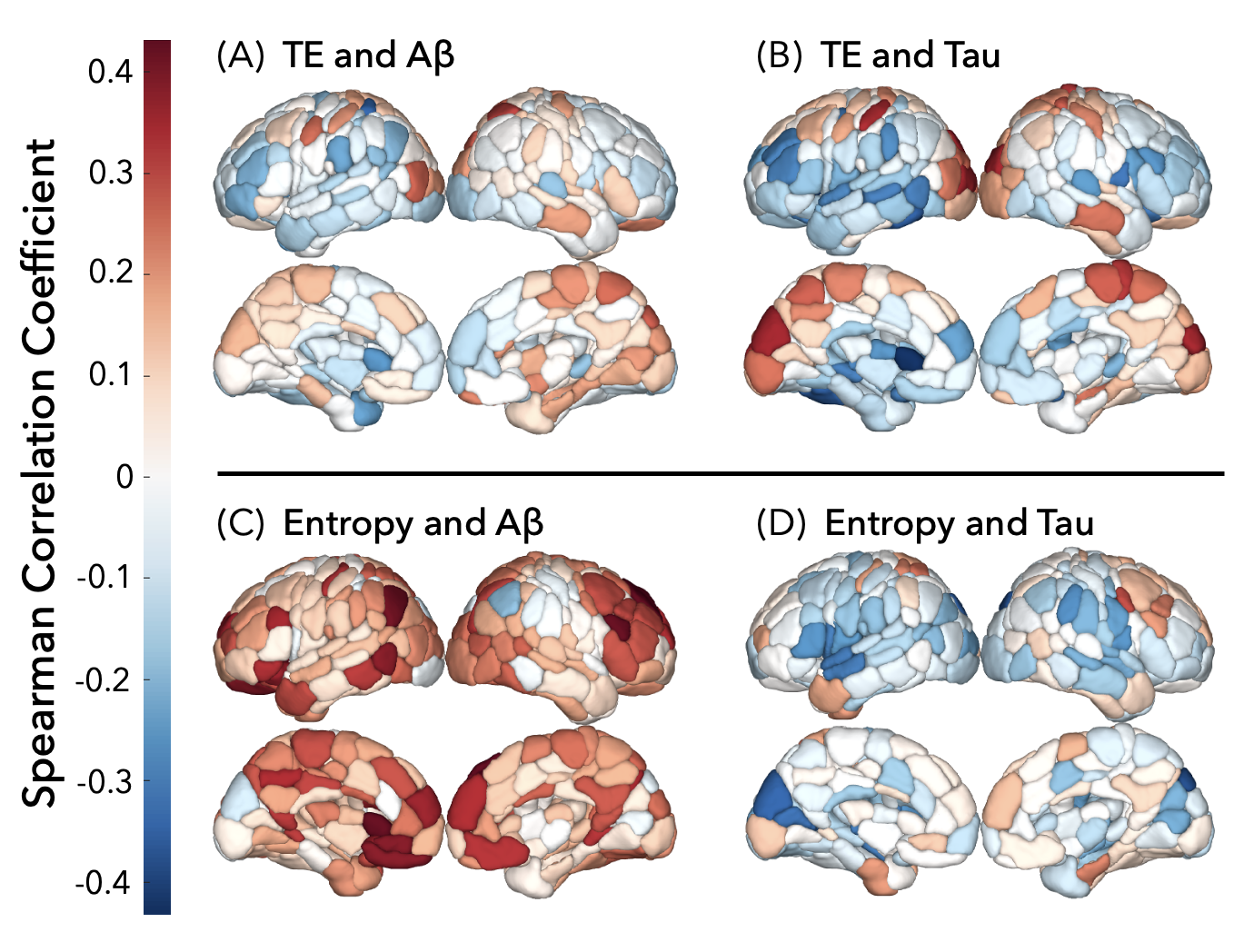

**Supplementary Figure 6**: Correlation between regional TE and PET metrics (Aβ and tau) in MCI patients. Spearman’s correlation coefficient between the regional (A) TE and Aβ, (B) TE and tau, (C) entropy and Aβ, and (D) entropy and tau was presented in the figure. Aβ plaque and tau protein levels are given in units of standard uptake value ratio (SUVR).

**Transition Energy Results Obtained with T = 1**

Supplementary Figure 7 shows the global, state-wise, network-wise, and regional TE in MCI and HC, as well as the beta estimates obtained from the linear model comparing these metrics between the MCI and HC groups. TE was calculated using T = 1. The linear model was adjusted for age, sex, and ICV. The reported *p*-values were also obtained from this linear model. Higher TE was observed in the visual and dorsal attention networks in both HC and MCI groups. Comparison revealed that 10 brain regions showed significantly higher TE in HC compared to MCI, and 29 regions showed significantly higher TE in MCI compared to HC after the multiple comparison *p*-value correction. The TE needed for the transition between the VIS– and DAN+ states was significantly higher in HC compared to MCI before correction, while the TE needed for transition between the VIS+ and DMN– states was significantly higher in MCI compared to HC before correction. Finally, the global TE was not significantly higher in HC compared to MCI (p = 0.901).

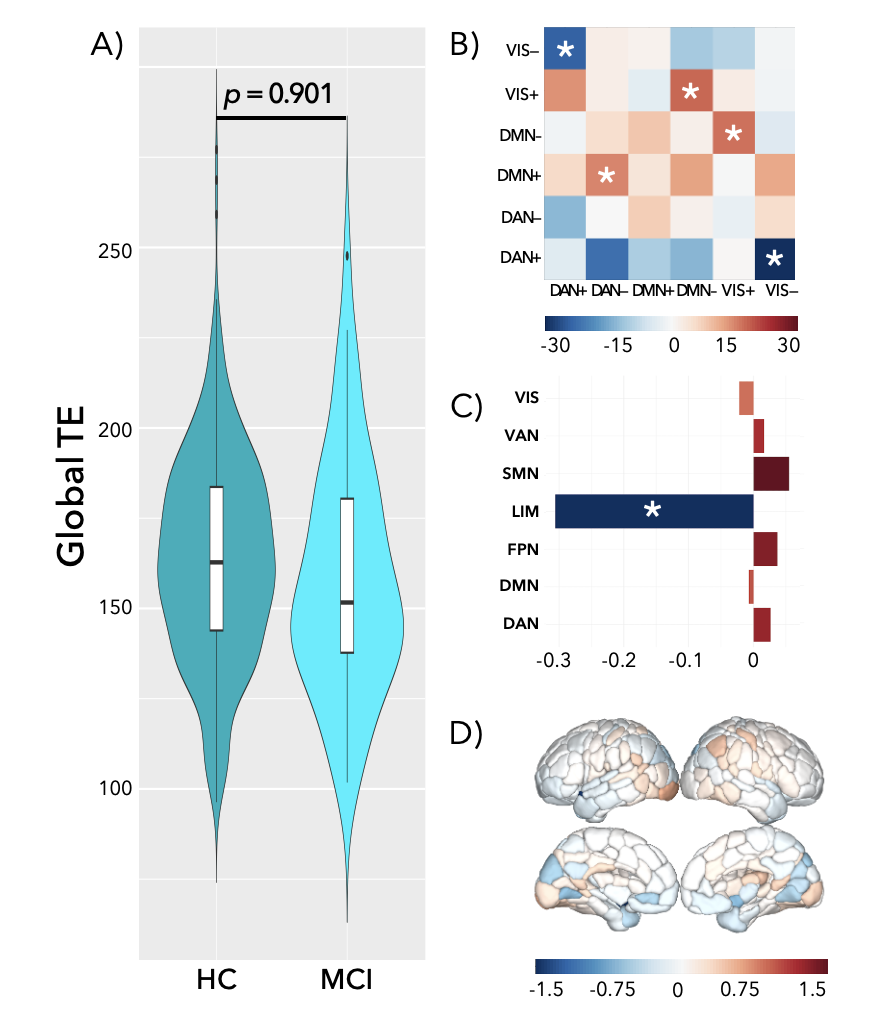

**Supplementary Figure 7**: (A) Distribution of global transition energy in HC and MCI. The p-value (p = 0.901) was computed using the linear model where the output was the global TE and the inputs were the group assignment (MCI vs. HC), age, sex, and ICV. (B) The beta estimates from the linear model adjusted for age, sex, and ICV were used to represent the amplitude and the direction of the difference in the state-wise TE between HC and MCI groups. State pairs with significantly different TE before correction are marked with an asterisk. None of the state pairs were significant after correction. (C) The beta estimates from the linear model adjusted for age, sex, and ICV were used to represent the amplitude and the direction of the difference in the network-wise TE between HC and MCI groups. The limbic network (marked with an asterisk) was significant after multiple comparison *p*-value correction. (D) The beta estimates from the linear model adjusted for age, sex, and ICV are used to represent the amplitude and the direction of the difference in the regional TE between HC and MCI groups. In figures B-D, positive values represent higher TE in MCI compared to HC, while negative values represent higher TE in HC compared to MCI group.

Supplementary Figure 8 is the replication of Figure 5 using T = 1 instead of T = 0.501 to generate the TE. The results are similar with T = 1 and T = 0.501.

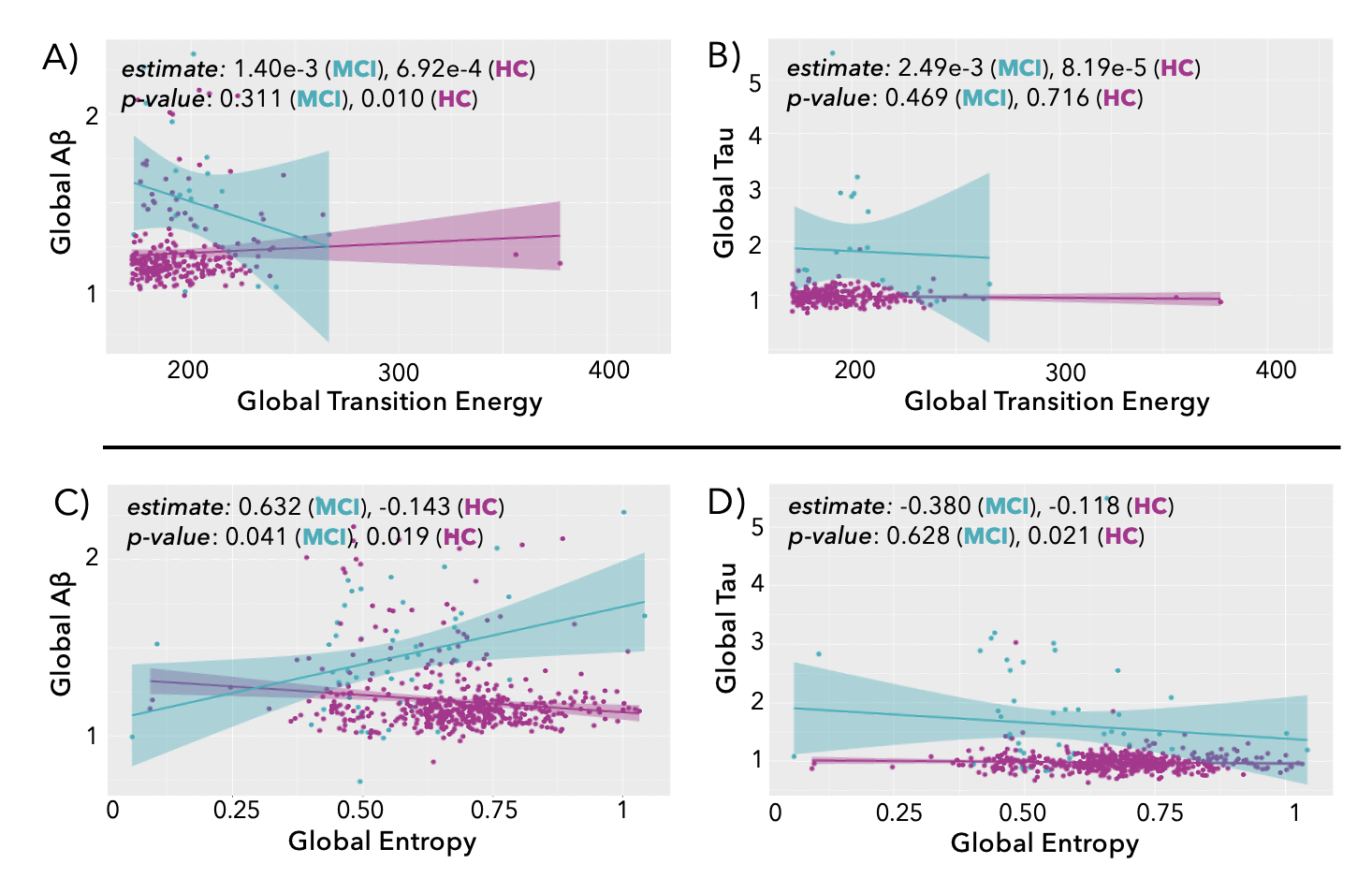

**Supplementary Figure 8**: The scatter plots represent the association between (A) global TE calculated with T = 1 and global entropy, (B) global TE calculated with T = 1 and global tau protein levels, (C) global TE calculated with T = 1 and global Aβ plaque levels, (D) global entropy and global tau protein levels, and (E) global entropy and global Aβ plaque levels. The data shown in this figure represents both HC and MCI patients. Aβ plaque and tau protein levels are given in units of standard uptake value ratio (SUVR). The β and the *p*-value from the linear model adjusted for age, sex, and ICV are presented on the scatter plots.
